# Addition of Genetics to Quantitative MRI Facilitates Earlier Prediction of Dementia: A Non-invasive Alternative to Amyloid Measures

**DOI:** 10.1101/731661

**Authors:** Natalie Marie Schenker-Ahmed, Nafisa Bulsara, Lei Yang, Lei Huang, Arya Iranmehr, Jian Wu, Alexander M Graff, Tetiana Dadakova, Hyun-Kyung Chung, Dmitry Tkach, Ilan Shomorony, Naisha Shah, Peter Garst, Robyn Heister, Svetlana Bureeva, Wayne Delport, David S Karow, James B Brewer, Christine Leon Swisher, for the Alzheimer’s Disease Neuroimaging Initiative

**Affiliations:** Human Longevity, Inc., San Diego, CA, USA; Department of Radiology, UC San Diego School of Medicine, La Jolla, CA; Department of Neurology, UC San Diego School of Medicine, La Jolla, CA

## Abstract

**Background:** Alzheimer’s disease is a major health problem, affecting ~4⋅5% of people aged 60 and older in 2016 with over 43 million affected globally^1^. The traditional approach for detection evaluates an individual in the presence of symptoms. However, it has been established that amyloid deposits begin to accumulate years before symptoms begin to appear^2,3^. With improved technology, there is increased focus on risk reduction, timely diagnosis, and early intervention. Early identification of at-risk individuals may enable patients and their families to better prepare for and reduce the impact of this condition.

**Methods:** We obtained data for patients from two longitudinal retrospective cohorts (Alzheimer’s Disease Neuroimaging Initiative: ADNI and National Alzheimer’s Coordinating Center: NACC), including T1-weighted MRI and genetics data. The polygenic risk score (PRS) used in this study was built based on a published Genome Wide Association Study (GWAS) that identified variants associated with Alzheimer’s disease. Quantitative MRI features were obtained using a 3D U-Net neural network for brain segmentation. Cox proportional hazards (CPH) regression models were used with subjects censored at death or the last evaluation. Time-to-event was defined as the time it takes for an individual who is dementia-free at the baseline MRI to progress to dementia as defined by the criteria described by ADNI. Time-dependent ROC areas under curve (AUCs) were estimated in the presence of censored data. The time-dependent AUCs were compared among models using the Wilcoxon rank sum test for dependent samples. Data was binned into three groups according to survival probability to eight years after baseline and Kaplan-Meier survival analysis was used to estimate the probability of surviving at least to time t. Calibration for both training and validation cohorts was evaluated using the predicted survival probability, splitting samples into five risk groups of equal size based on the predicted survival probability.

**Findings:** We developed a model that predicts the onset of dementia over an eight-year time window in individuals with genetics data and a T1-weighted MRI who were dementia-free at baseline. We then validated the model in an independent multisite cohort.

We observed that models using PRS in addition to MRI-derived features performed significantly better as measured by time-varying AUC up to eight years in both the training (p = 0⋅0071) and validation (p = 0⋅050) cohorts. We observed improved performance of the two modalities versus MRI alone when compared with more invasive amyloid measures. The combined MRI and PRS model showed equivalent performance to cerebral spinal fluid (CSF) amyloid measurement up to eight years prior to disease onset (p = 0⋅181) and while the MRI only model performed worse (p = 0⋅040). Finally, we compared to amyloid positron emission tomography (PET) three to four years prior to disease onset with favorable results.

**Interpretation:** Our finding suggests that the two modalities are complementary measures, in that MRI reflects near-term decline and the addition of genetics extends the prediction scope of quantitative MRI by adding additional long-term predictive power.

The proposed multimodal model shows potential as an alternate solution for early risk assessment given the concordance with CSF amyloid and amyloid PET. Future work will include further comparison with amyloid PET (greater than four years) and with CSF (greater than eight years) as additional long-term data becomes available. Also, the model will be evaluated for its clinical utility in the “active surveillance” of individuals who may be concerned about their risk of developing dementia but are not yet eligible for assessment by amyloid PET or CSF.

**RESEARCH IN CONTEXT:** 

**Evidence before this study:** The most significant known genetic factor in Alzheimer’s disease (AD) is the ε4 allele for the *Apolipoprotein E* (*APOE*) gene. Carriers of the allele have a three-fold increased risk of developing AD, whereas individuals who are homozygous have a 15-fold increased risk. Genome-wide association studies (GWASs) have identified many additional genetic variants that are associated with AD. Recent studies have shown that the risk for AD is better predicted by combining effects from several genetic variants into “polygenic risk scores” (PRS). Studies have also demonstrated that the age of onset for AD is better predicted using PRS rather than *APOE* status alone. Regional brain atrophy, as measured using volumetric MRI, is also an important biomarker for evaluating an individual’s risk of developing dementia. Previous predictions have shown that medial temporal lobe atrophy, as measured by a Hippocampal Occupancy Score (HOC) is highly associated with progression from MCI to AD.

**Added value of this study:** In the proposed model, the addition of genetics to MRI data lengthens the time over which the model can predict onset of dementia. The two measures appear to be complementary, with MRI showing near-term decline and genetics providing additional predictive power in the long-term. When compared to more invasive measures of amyloid, which have been shown to have long-term predictive power, we observed equivalent performance to CSF amyloid up to 8 years prior to disease onset and equivalent performance to amyloid PET three to four years prior to disease onset.

**Implications of all the available evidence:** Although MRI remains relatively expensive, it is less expensive, less invasive, more accessible, and more commonly available than amyloid PET. Furthermore, MRI is already part of standard clinical practice and this model may be applied to standard clinical MRIs with no additional acquisition required. A recent survey of patients and their caregivers has highlighted a desire for access to better diagnostics, such as amyloid PET, to aid them in long-term legal, financial and healthcare planning. Our model, given the concordance with CSF and amyloid PET could be an alternate solution to fulfill this need. Furthermore, our model could facilitate the “active surveillance” of individuals who are high-risk and thereby enhance the possibility of early intervention.

## INTRODUCTION

### Description of Problem and Motivation

Dementia is a clinical syndrome characterized by progressive deterioration in cognitive ability and reduction in capacity for independent living and functioning that results from brain damage. Brain damage can occur due to a variety of causes with the most common being Alzheimer’s disease (AD), although many Alzheimer’s patients also have damage resulting from vascular disease. Dementia is a major global health problem, affecting ~4⋅5% of people aged 60 and older in 2016 with over 43 million affected globally^1^. It exacts tremendous costs, both in terms of disability for those affected and in pure economic terms. The predicted cost of dementia is greater than $290 billion yearly in the United States alone^4^, and the disease is responsible for 30 million disability adjusted life years^5^. In the absence of a cure and with improved technology, there is increased focus on risk reduction, timely diagnosis, and early intervention. Early identification of at-risk individuals may enable patients and their families to better prepare for and reduce the impact of this condition.

The traditional approach evaluates an individual in the presence of symptoms. However, it has been established that amyloid deposits begin to accumulate years before symptoms begin to appear^2,3^. Furthermore, lifestyle is key in influencing an individual’s risk^6^, and early risk identification provides greater opportunity for intervention and risk mitigation^7^. To accomplish the early identification of at-risk individuals, prediction must rely on evidence prior to the onset of symptoms. In certain diseases, such as coronary artery disease, there are clear biomarkers and calculated scores that aid physicians in predicting which patients are at high-risk. Although significant research effort has focused on the development of better models for predicting onset of dementia, particularly in the progression of individuals from mild cognitive impairment (MCI) to AD^8–10^, a predictive score, used clinically, that assess and individual’s risk is yet to be developed. Biomarkers that may be used include genotype, in particular *APOE* status, and quantitative magnetic resonance imaging (MRI). Testing for amyloid protein in the cerebral spinal fluid or in the brain may also be performed to determine whether a diagnosis of Alzheimer’s is indicated^11^.

### *APOE* genotype and other genetic risk analyses

The most significant known genetic factor in AD is the ε4 allele for the *Apolipoprotein E* (*APOE*) gene. Carriers of the allele (heterozygous state with one of the other common variants) have a three-fold increased risk of developing AD, whereas individuals who are homozygous have a 15-fold increased risk. 20-25% of individuals with AD have at least one copy of the ε4 allele^12^. However, late onset dementia is a complex disorder influenced by environmental and genetic factors, with *APOE* variants accounting for the majority, but not all, of phenotypic variance^13^. To this end, genome-wide association studies (GWASs) have facilitated the identification of additional genetic variants associated with AD, although most have a limited effect^13–15^ and novel associations continue to be uncovered^16^. Some researchers have combined the effects of known variants into a polygenic hazard score. Such combined genetics scores have demonstrated better prediction of the age of onset for AD versus *APOE* status alone^15,17^ and may correlate with amyloid beta and tau accumulation, and cortical volume changes^17^.

### Quantitative MRI

Regional brain atrophy, as measured using volumetric MRI, is another important biomarker for evaluating an individual’s risk of developing dementia. Previous predictions have shown that medial temporal lobe atrophy, as measured by a Hippocampal Occupancy Score (HOC) is highly associated with progression from MCI to AD^18,19^. Analysis of the signal intensity of the hippocampus and comparison to cognitively normal brains vs Alzheimer’s may also catch early evidence of Alzheimer’s disease^20^.

### Amyloid ß and Tau testing

Presence of Alzheimer’s disease is also evaluated by determining whether significant levels of amyloid beta (Aß) proteins^21^ or tau^22^ are present in the CSF or in the brain. Results are acquired either via lumbar puncture or amyloid positron emission tomography (PET) imaging. Regardless of the method, measurement of Aß deposition in the brain may be used to stratify patients into high and low risk categories, and may impact clinical management^21,23^.

Further studies using PET imaging have led to assessment of its utility for predicting disease progression using machine learning models. One such algorithm demonstrated the utility of using the information from a single PET scan in the prediction of disease progression for MCI individuals within 2 years^24^.

Testing for Aß or tau, either via lumbar puncture or PET imaging is invasive (because of the need for an intravenous injection of radioactive dye), although work is progressing toward a less invasive blood test for tau^25^. Furthermore, although such testing may be available to the research community, it is not readily accessible in a clinical setting, due to low availability and high expense. As a result, it is neither appropriate nor feasible to perform testing for Aß or tau as a screening procedure or risk assessment for asymptomatic individuals.

### Prediction models in the literature

To our knowledge, no existing model has integrated data to predict progression to dementia by cognitive normal individuals many years prior to diagnosis. Most existing models rely heavily on cognitive test results to drive their prediction and many have focused on the progression of individuals with MCI^26,27^. Predictions have been made using genetics alone^8^, MRI alone^9,28^, and through the combination of genetics with MRI and cognitive testing^10,15,17^.

### Modeling disease risk with Survival Models

Survival Analysis is a statistical framework used to analyze how a set of variables influence a subject’s survival time or, more generally, the time to an event of interest. Typically, the goal of the analysis is to build a regression model to predict the time to event for each subject based on a set of covariates. A common challenge in building such models is that a fraction of the data is often censored, that is, the subject leaves the study before the occurrence of the event of interest.

The Cox proportional hazards (CPH) model is a standard statistical model for investigating the association between survival time and predictor variables and aims to compute a hazard function for each individual, which describes how the risk of the event evolves with time. The proportional hazards model assumes that the hazard function consists of two parts: a baseline hazard function, which is common to the whole population, and a multiplicative factor, which is unique for each individual.

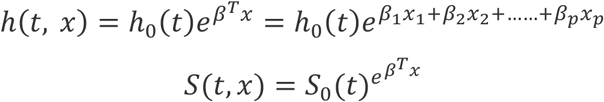

Where *h*(*t*, *x*) is the hazard function for people at risk with predictors *x*, *h*_0_(*t*) is the baseline hazard function for the population of people with *x=0*, *S*(*t*, *x*) is the individual survival function, i.e., the probability a person’s event occurs after each time t, and *S*_0_(*t*) is the baseline survival function (*x=0*). A powerful property of the CPH model is that it can handle censored data.

### Proposed Solution

The gold standard for diagnosis of Alzheimer’s disease remains post-mortem neuropathological inspection, but amyloid PET has been found to be a useful diagnostic tool for Alzheimer’s disease in living patients^29^. Successful tools to identify Alzheimer’s disease earlier include those based on the presence of characteristic proteins in the CSF or in the brain. However, the detection of Aß via PET or lumbar puncture has significant drawbacks, limiting the clinical utility for low risk and asymptomatic individuals. Although MRI remains relatively expensive, it is less expensive, more accessible, and more commonly available than amyloid PET. Therefore, a model for prediction of dementia onset using MRI and genetics, rather than the more expensive and more invasive Aß testing, would make early prediction accessible for many more individuals. In the current study, we developed an improved prediction model leveraging quantitative MRI features and genetics to assess individuals’ long-term risk of dementia (Figure 1).

**Figure 1:**
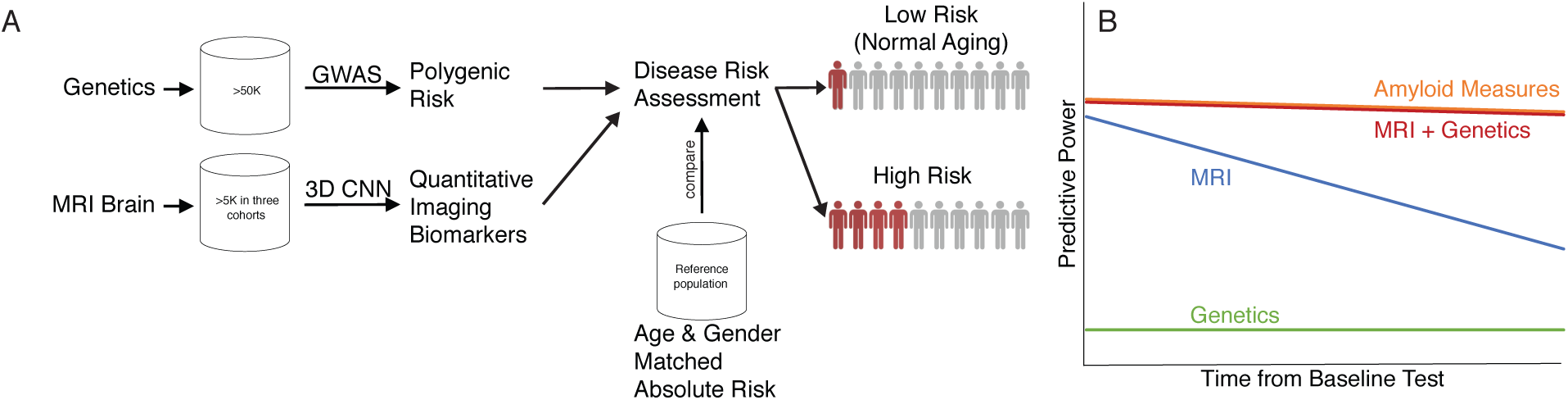
Overview of the proposed model. A) The model uses inputs of genetics data and MRI data to determine an individual’s absolute risk of dementia. Age and gender matched absolute risk comparisons with a reference cohort could be used to stratify low risk and normal aging from high risk individuals. B) The schematic describes the proposed model’s expected performance for each input modality. We hypothesize that combining MRI and genetics data is expected to have similar performance to amyloid measures and have improved performance than either MRI or genetics alone.

Our model for an individual’s risk of developing Alzheimer’s primarily uses data from two sources: 1) the individual’s genotype, specifically regarding the gene variants that are known to be associated with an inherited risk of developing Alzheimer’s and 2) an individual’s current neuroanatomical phenotype, using structural measurements derived from fully convolutional neural networks from MRIs of their brain.

## MATERIALS

### Training Cohort

All longitudinal 3T and 1⋅5 T brain MRIs along with SNP genotype data, associated baseline diagnosis and demographic information were obtained from the Alzheimer’s Disease Neuroimaging Initiative (ADNI) database (adni.loni.usc.edu). The ADNI was launched in 2003 with the primary goal of ADNI of testing whether serial magnetic resonance imaging (MRI), positron emission tomography (PET), other biological markers, and clinical and neuropsychological assessment can be combined to measure the progression of AD. For up-to-date information, see www.adni-info.org.

### Independent Multisite Validation Cohort

The genetic data for this study were prepared, archived, and distributed by the National Institute on Aging Alzheimer’s Disease Data Storage Site (NIAGADS) at the University of Pennsylvania, and the baseline MRI was obtained from National Alzheimer’s Coordinating Center (NACC). NACC was established by the NIA/NIH to facilitate collaborative research between Alzheimer’s Disease Centers (ADC). The baseline MRI data used in this study was collected since 2005 as a part of NACC Uniform Data Set (UDS)^30^. Eleven individuals known to exist in both cohorts were excluded from the validation cohort.

## METHODS

### MRI features

Quantitative MRI features were obtained from 3D brain MR images using a deep learning neural network method for brain segmentation^31^. This fully automated method improves upon existing semi-automated methods of brain segmentation by being very fast and consistently reproducible. Measurements of amygdala and hippocampus volume were included in the prediction model directly. For a subset of the validation cohort, MRI features were pulled from previously extracted features in the validation database, as the MR images were not available for processing with the above-mentioned method.

### Polygenic Risk Score

The polygenic risk model used in this test was built based on a published Genome Wide Association Study (GWAS) that identified variants associated with Alzheimer’s disease (Supplemental Table S1)^14^. This GWAS was performed in individuals of European ancestry comprising approximately 26,000 cases and 48,000 controls. Additionally, the model includes the *APOE* ε4 variant, a well-known risk factor for Alzheimer’s disease^32^. The GWAS approach allows for the estimation of a variant-specific weight that is directly proportional to the strength of the association between the variant and the disease. A polygenic risk score is the sum of these weights for all variants that are shared between the polygenic risk model and the genome of the subject.

### Survival Model training

Cox proportional hazards (CPH) regression models were used with subjects censored at death or the last evaluation. In the clinical setting, a CPH model can be used as a prediction tool to estimate an individual’s relative and absolute risk of developing disease at time t. Survival models have the advantage of accounting for the variable duration of follow-up, time-to-event and censoring. Time-to-event was defined as the time it takes for an individual who is dementia-free at the time of the baseline MRI to progress to dementia as defined by the following criteria described by ADNI: Memory complaint by subject or study partner; Abnormal memory function by education adjusted cutoff on the logical memory ii subscale; Mini-mental state exam score (MMSE) between 20-26; Clinical dementia rating = 0⋅5 or 1⋅0; NINCDS/ADRDA criteria for probable AD. The baseline measurement was the date of the first MRI in the study. We leveraged the Breslow method^33,34^ to handle ties, where failures are only reported with day accuracy. At baseline, all individuals were dementia free, with a mixture of individuals with normal and MCI classification. Conversion rate was 12⋅5% in the internal training cohort and 19⋅3% in the external validation cohort.

### Survival Model Features

Features were selected based on a significant p-value (<0⋅05) and feature importance using Cox (Table 2). A derived feature, HOC, was calculated as indicated in Table 3 and normalized using a natural log transformation. The contribution of genetics to risk was assessed with the inclusion of polygenic risk as calculated according to values presented in Supplemental Table S1. All feature processing steps were the same on the external validation cohort as the internal training cohort.

**Table 1:**
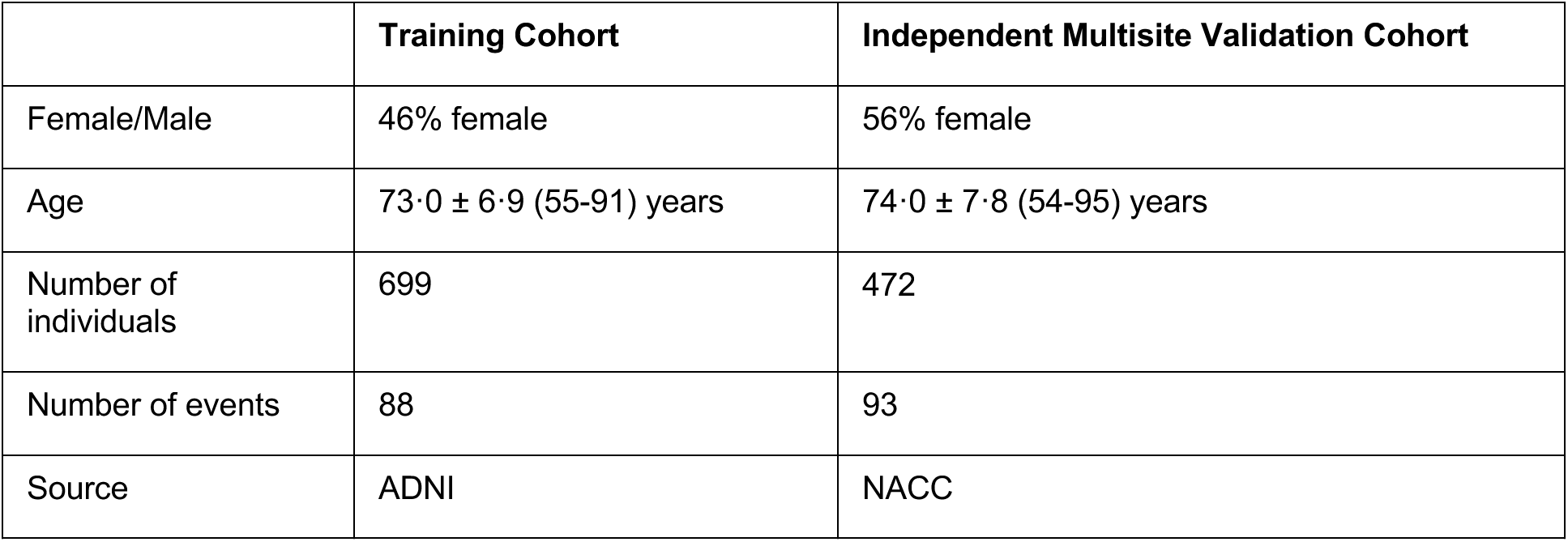
Cohort description

**Table 2:** Features included in the integrated risk model for dementia

|  | Feature Type | Definition | Feature Transformation Applied |
| --- | --- | --- | --- |
| Age | Demographic | Age at visit | Binned into two groups ($\leq 73.2$, $> 73.2$) |
| Amygdala volume | Imaging | Sum of left and right hemisphere volumes of the amygdalae - roughly almond-shaped masses of gray matter inside each cerebral hemisphere, involved with the experiencing of emotions. | Z-score transform |
| Hippocampal Occupancy Score (HOC) | Imaging | $HOC = H / (ILV + H)$ where:<br>ILV = Sum of left and right hemisphere volumes of the inferior lateral ventricles. | Natural logarithm, then Z-score transformation |
|  |  | H = Sum of left and right hemisphere volume of the hippocampi |  |
| GWAS | Genetics | Polygenic risk score computed as defined above. | Z-score transform |

Since brain features are highly correlated with age, we leveraged a stratified model based on age rather than leveraging age as an input feature. We used only two age bins to prevent overfitting. When using the stratified model, the baseline hazards are allowed to be different for different strata. The hazard for an individual from stratum k is given by:

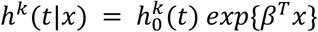

where 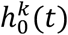 is the baseline hazard for stratum *k*, *k* = 1, …, *k*

### Model discrimination performance and generalization

We evaluated the model’s ability to discriminate risk at each year using inverse probability censored weighting (IPCW) to measure the time-varying area under the curve (AUC) from receiver operating characteristic (ROC) curves and overall performance with concordance index. In the context of survival modeling, concordance index is the probability that for a pair of randomly selected samples, the sample with larger predicted risk will experience an event before the other. Details of the models and the relevant C-indices can be found in Supplemental Table S2.

Time-dependent ROC curves and AUCs were estimated in the presence of censored data (Figure 2A&B). Time-varying AUCs for the MRI only, MRI and Genetics, CSF, and Amyloid PET models were compared using a Wilcoxon rank sum test for dependent samples. Five-fold cross validation was used to assess model performance on the training cohort. A test and train split strategy was not used during training due to a small data set.

**Figure 2:**
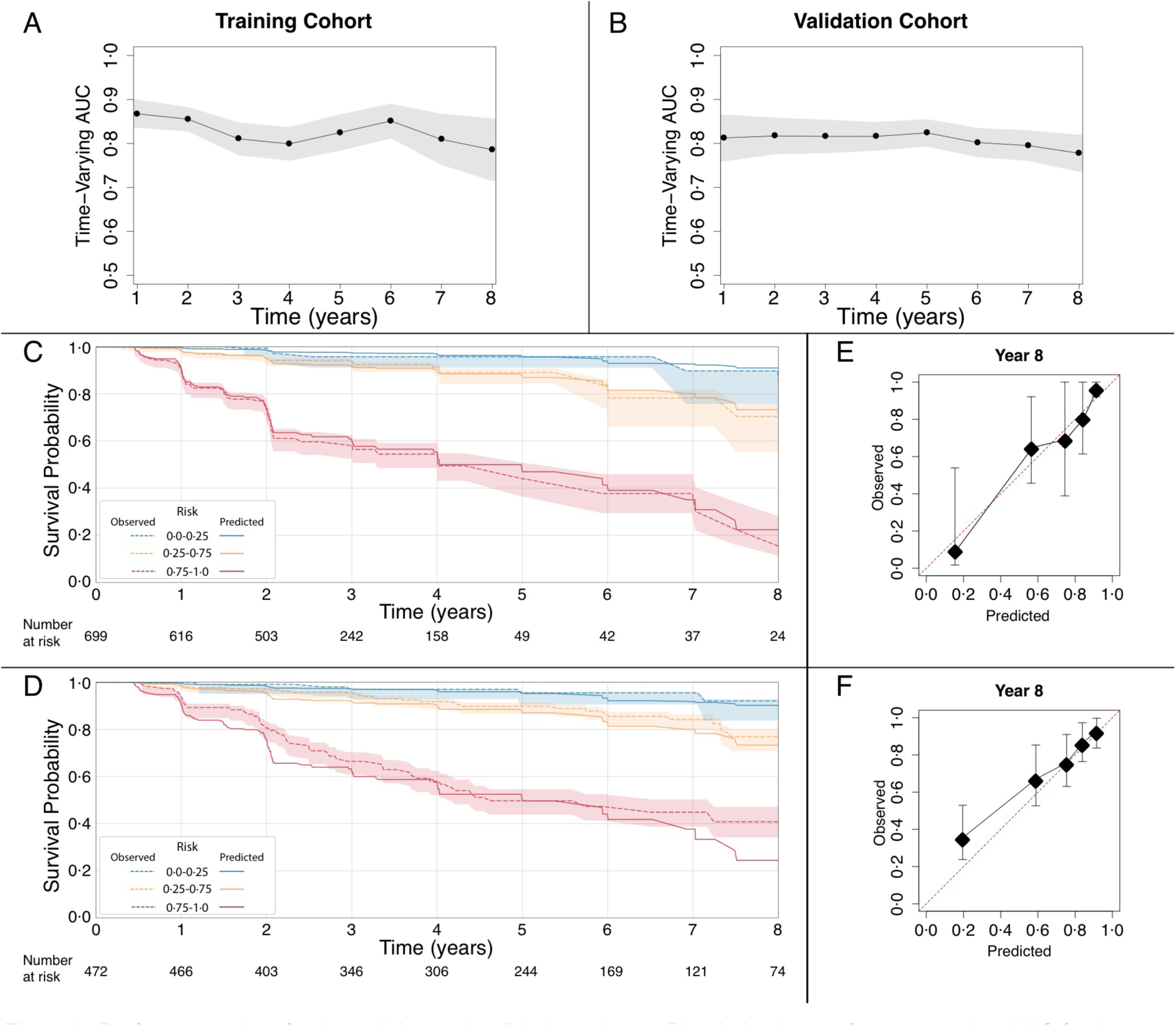
Performance data for the training and validation cohorts. Discrimination performance using AUC for the ROC curves at years 1-8 prior to a progression event in the A) training and B) validation cohorts. Kaplan Meier plots showing survival probability from baseline to 8 years in the C) training and D) validation cohorts. Data is binned by predicted risk of developing dementia within 8 years from baseline: bin1: 0⋅0-0⋅25; bin 2: 0⋅25-0⋅75; bin3: 0⋅75-1⋅0) in the E) training and F) validation cohorts. Calibration plots for prediction 8 years prior to a progression event.

### Calibration

Both discrimination and calibration are critical for model validation. Even if a new prediction model discriminates well, if good calibration is not also achieved the model may perform poorly in a new patient population. Despite this, calibration is rarely reported in risk prediction studies^35^. We tested the combined MRI and Genetics model’s calibration, or its ability to produce unbiased estimates of risk, by measuring the degree of agreement between the estimated and observed survival probability. Survival probability estimations^33,34^ were collected for time periods from one year from baseline through the length of the testing cohort study (eight years).

These calibration curves were visually inspected to determine whether or not the observed event frequencies matched the expected event probabilities for each group of samples. Calibration curves were expected to produce a linear relationship between observed and expected probability of dementia and not deviate strongly from a 1:1 relationship. Plots were created for calibration in both the training (ADNI) and test (NACC) datasets, respectively (Figure 2D&E, Supplemental Figure S1). The survival probability was predicted using the model built on training dataset (x-axis). All samples were split into five risk groups of equal size based on the predicted survival probability. The observed survival probability was estimated using Kaplan-Meier estimator on each risk group (y-axis).

The strong agreement of predicted versus observed risk is expected for the cohort on which the model was trained (Figure 2E at eight years; Supplemental Figure S1A for years two, four, and six). The degree of over- or under-estimation of risk is minimal. Similarly, there is limited over- and under-estimation of risk in the validation cohort (Figure 2F at 8 years; Supplemental Figure S1B for years two, four, and six).

### Survival analysis

Kaplan-Meier survival analysis was used to estimate the probability of surviving at least to time t (Figure 2C&D). Data was binned according to the predicted risk of developing dementia within 8 years from baseline in the following three risk groups: probability = 0⋅0-0⋅25 and 0⋅25-0⋅75 and 0⋅75-1⋅0 and compared to the actual survival curve for the individuals in each bin. Error shown is standard deviation. We observe significant separation of the three groups (p<0⋅0001 for both training and validation cohorts) as well as good concordance of the observed and predicted survival curves.

## Results

### PRS vs *APOE*

A model using *APOE* alone instead of the PRS was built on the training dataset and evaluated for predictive performance in comparison with the one using the PRS. Figure 3 illustrates the concordance index for each model in both the training and validation cohorts. In both cohorts, PRS produces a significantly higher C-index (Wilcoxon rank sum test). Statistics of the different genetic models are shown in Supplemental Table S3, in which the effect sizes for PRS are reported as the hazard-ratio per standard deviation.

**Figure 3.**
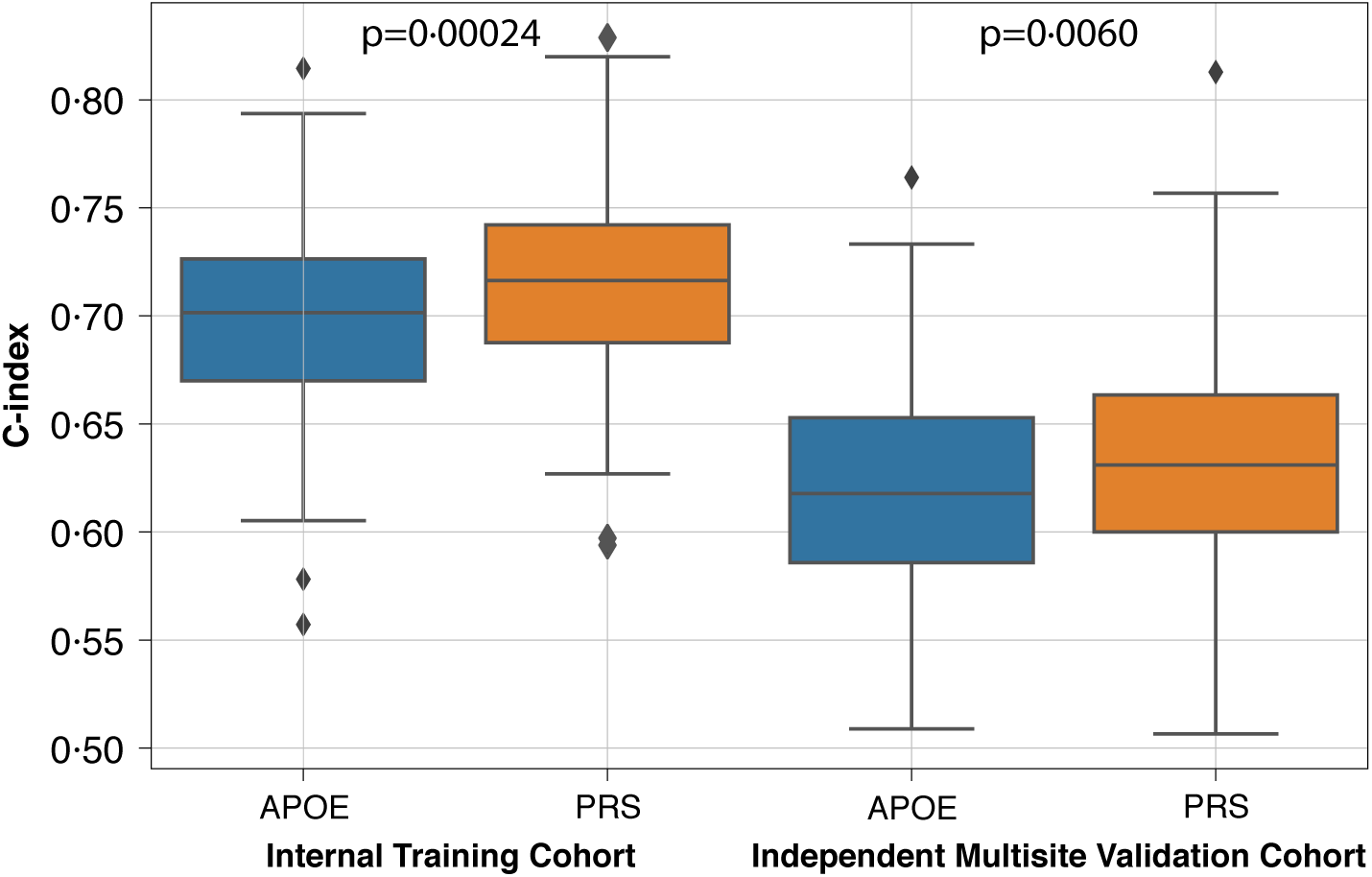
Comparison of models using PRS versus APOE alone. C-indices were significantly higher in both the training (p=0.00024) and validation (p=0.0060) cohorts.

### MRI/PRS vs MRI only

Two models (MRI only and combined MRI/PRS) were built on the training dataset and time-varying AUC was computed on both training and test datasets using the two models respectively (Figure 4). The AUCs in the combined model were significantly greater than those in the MRI only model in the training cohort (p=0⋅0071). Also, the C-indices for were higher for the combined model vs MRI only in both the training (MRI/PRS CI=0⋅82; MRI CI=0⋅79) and validation (MRI/PRS CI=0⋅807, MRI CI=0⋅788) datasets.

**Figure 4:**
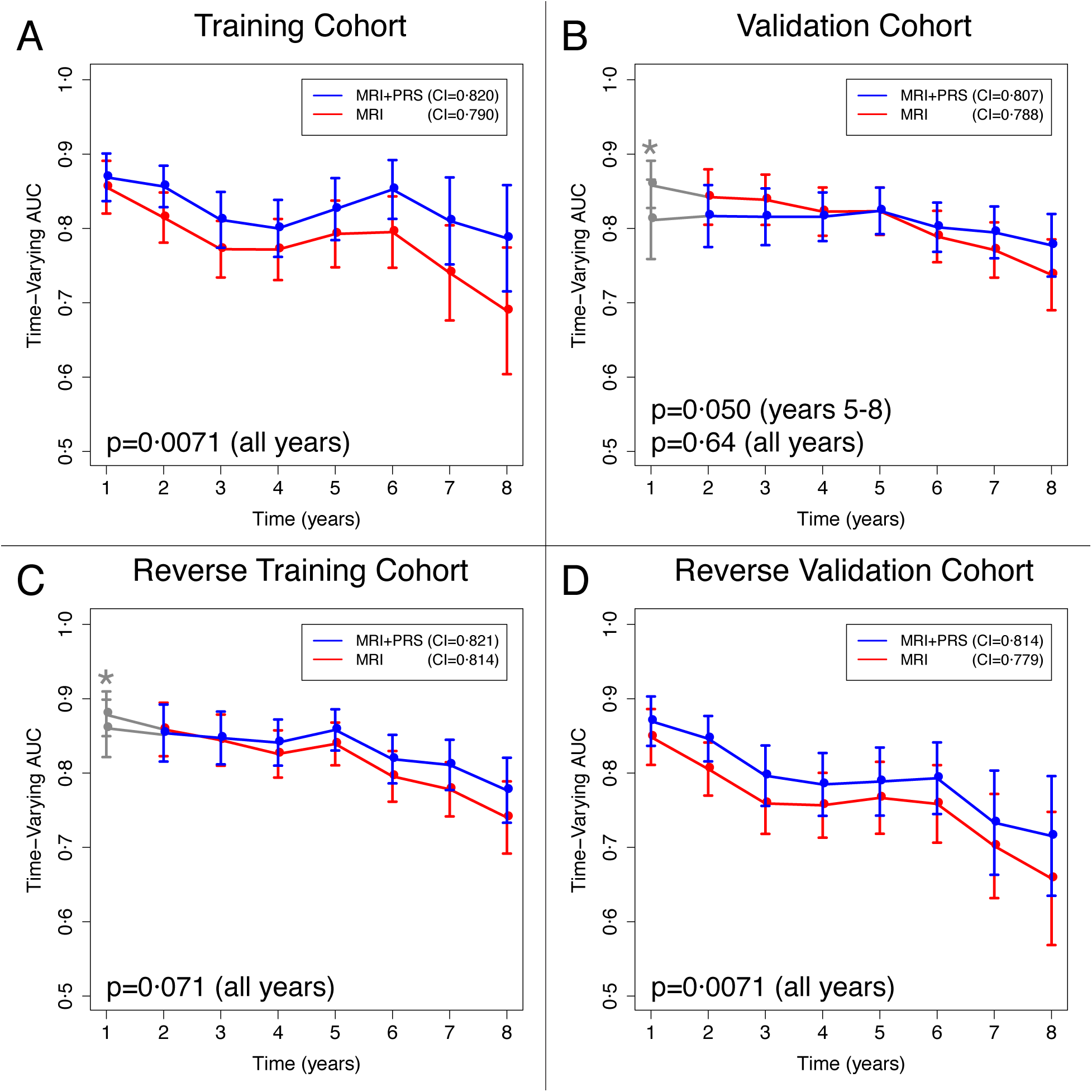
Comparison of the proposed model using MRI and PRS versus a model using only MRI features in the A) training and B) validation cohorts. A similar trend is seen in the comparison of time-varying AUC between MRI+PRS and MRI alone when the training and validation cohorts were reversed (C&D). *In the validation cohort (and therefore, the reverse training cohort), there were few events at year one in the validation cohort and no events in high genetic risk individuals within the first year (see Supplemental Figure S2 and Table S4).

### Comparison with Amyloid Beta

Models were built on the training dataset using CSF amyloid levels or amyloid PET (florabetapir) and time-varying AUC was computed on both training and test datasets for comparison with the combined model (Figure 5). MRI/PRS showed equivalent performance to cerebral spinal fluid (CSF) amyloid measurement up to eight years prior to disease onset (p = 0⋅181), whereas MRI alone showed worse performance (p = 0⋅040).

**Figure 5:**
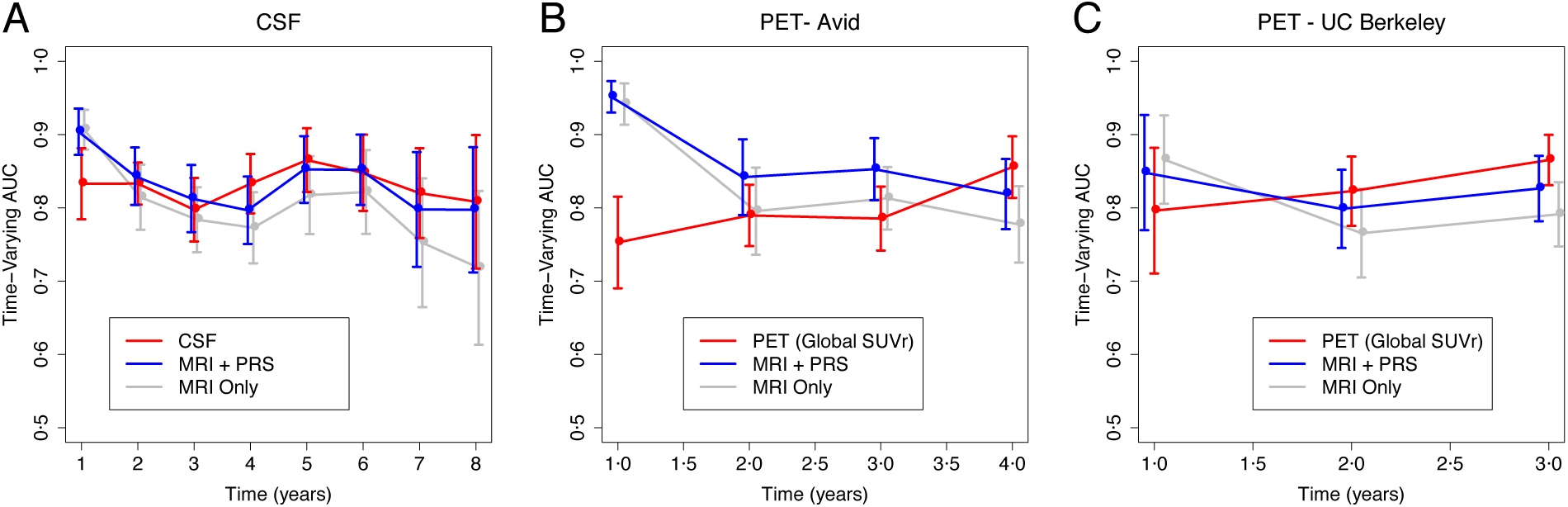
Comparison of MRI/PRS model, MRI only model, and A) a model using CSF amyloid levels, B) a model using Global SUVr in an Avid study^36^; C) a model using Global SUVr in a UC Berkeley study^37^.

## Discussion

Individuals who are asymptomatic but are concerned about their risk of dementia currently have few tools to assess their potential risk. Widely available genetic testing, such as *APOE* status, may result in a false sense of security, or cause undue alarm. More reliable tests, such as the detection of Aß via PET or lumbar puncture have significant drawbacks, limiting the clinical utility for low risk and pre-symptomatic individuals. Nonetheless, early risk identification is desirable, as it provides greater opportunity for intervention and risk mitigation^7^, potentially enabling individuals and families to better prepare for, postpone the effects of and reduce the severity of the disease’s impacts. In particular, modifiable lifestyle factors have a significant influence on an individual’s risk^6^, including key factors such as management of type II diabetes, managing hearing loss, and reducing alcohol use^38–40^.

Although MRI remains relatively expensive, it is more affordable, more accessible, and more commonly available than amyloid PET. Therefore, a model for prediction of dementia onset using MRI and genetics, rather than the more expensive and more invasive Aß testing, would make early prediction available for many more individuals.

In this work, we sought to develop a method that could be accessible to low-risk and pre-symptomatic individuals.

### Significant findings

We show that the integration of polygenic risk with deep learning-derived structural MRI features improves the prediction of dementia onset. While structural MRI is a strong biomarker by itself for both detection and short-term prediction, it does not have the ability to differentiate individuals prior to the anatomical changes which occur later in the disease trajectory. The addition of genetics in the form of a PRS to quantitative MRI features extends the time at which a model can accurately predict progression to dementia (Figure 4).

We hypothesize that the two modalities are complementary measures. MRI reflects near term decline while the genetics extends the prediction scope of quantitative MRI by adding long-term predictive power. Support of this hypothesis is provided by the improved performance of the combined model versus MRI alone when compared with more invasive amyloid measures (Figure 5). Furthermore, we show that the use of structural MRI features in combination with polygenic risk has equivalent performance to amyloid CSF up to eight years prior to disease onset (Figure 5A; p = 0⋅181; see Supplemental Table S2). We also see equivalent performance to amyloid PET 3-4 years prior to disease onset (Figure 5B&C; see Supplemental Table S2).

### Clinical application

Although MRI remains relatively expensive, it is less expensive, more accessible, and more commonly available than amyloid PET. Furthermore, MRI is already part of standard clinical practice, and the model proposed herein may be applied to standard clinical MRIs with no additional acquisition required. A recent survey of patients and their caregivers has highlighted a desire for access to better diagnostics, such as amyloid PET, to aid them long-term legal, financial and healthcare planning^41^. Given its concordance with CSF and amyloid PET, the proposed multimodal model shows potential as an alternate solution for early risk assessment. Conceivably, the model could be considered for active surveillance of individuals who are concerned about their risk of developing dementia but are not eligible for invasive testing such as amyloid PET.

### Additional applications

Since genetic sequencing and MRI are minimally invasive, our model could be used as a research subject risk stratification approach for clinical studies. For instance, identifying high-risk individuals would greatly enhance the efficiency of studies that are investigating the efficacy of new therapies for dementia.

### Limitations

Although, the *APOE* ε4 allele is a major risk factor for AD, odds ratios vary considerably among ethnic groups^42,43^, with the highest risk in East Asians^42,44^, followed by non-Hispanic Caucasians^42,45,46^ and lowest in African ancestry populations^42,47–49^. This may be the result of environmental factors, or population-specific genetic factors, with some evidence suggesting the latter^43^. Nonetheless, the odds-ratios derived from European studies of AD cannot be applied directly to non-European populations. Similarly, European-derived polygenic risk scores are not portable to other populations^50^. Although the current model is only applicable to Europeans, the polygenic risk score feature can easily be substituted with an ancestry-appropriate GWAS for application to non-European populations.

### Future work

We plan to investigate our model in application to larger and more diverse cohorts, as there were a limited number of events in our cohort. This may also allow us to improve our PRS model.

Also, further comparison with amyloid PET (greater than four years) and with CSF (greater than eight years) as additional long-term data becomes available. Also, the model will be evaluated for its clinical utility in the active surveillance of individuals who may be concerned about their risk of developing dementia but are not yet eligible for more invasive imaging such as amyloid PET.

## Conclusion

Early identification of individuals that are at-risk for dementia requires biomarkers that are evident prior to the onset of symptoms. Unlike many chronic conditions, there is no reliable risk score or assessment tool to help physicians identify high-risk patients. Other than standard cognitive tests, such as the MMSE, and basic genotyping, such as for APOE status, no simple test exists which can help individuals and their physicians assess an individual’s risk of developing dementia in the absence of symptoms.

Although MRI remains relatively expensive, it is already part of standard clinical practice, and genotyping is rapidly becoming more affordable and available. Therefore, the testing required for the model described in this manuscript is accessible and can be performed with minimal invasiveness, yet predicts with the same accuracy as the more invasive and less available CSF or amyloid PET. This model could greatly impact dementia screening. Early identification of high-risk individuals could facilitate the creation of “active surveillance” programs which could monitor individuals who are not yet symptomatic and assist them in making lifestyle changes which might postpone onset of symptoms or reduce their severity. Furthermore, active surveillance would also help individuals and their families to better prepare for the impacts of the disease.

## Acknowledgements

We thank Brad Perkins, Ying-Chen Claire Hou, and Keegan Duchicela for useful discussions. We thank Palak Sheth for data engineering support.

## Funding

Human Longevity, Inc

Alzheimer’s Disease Neuroimaging Initiative National Alzheimer’s Coordinating Center

---

**Table S 1.**
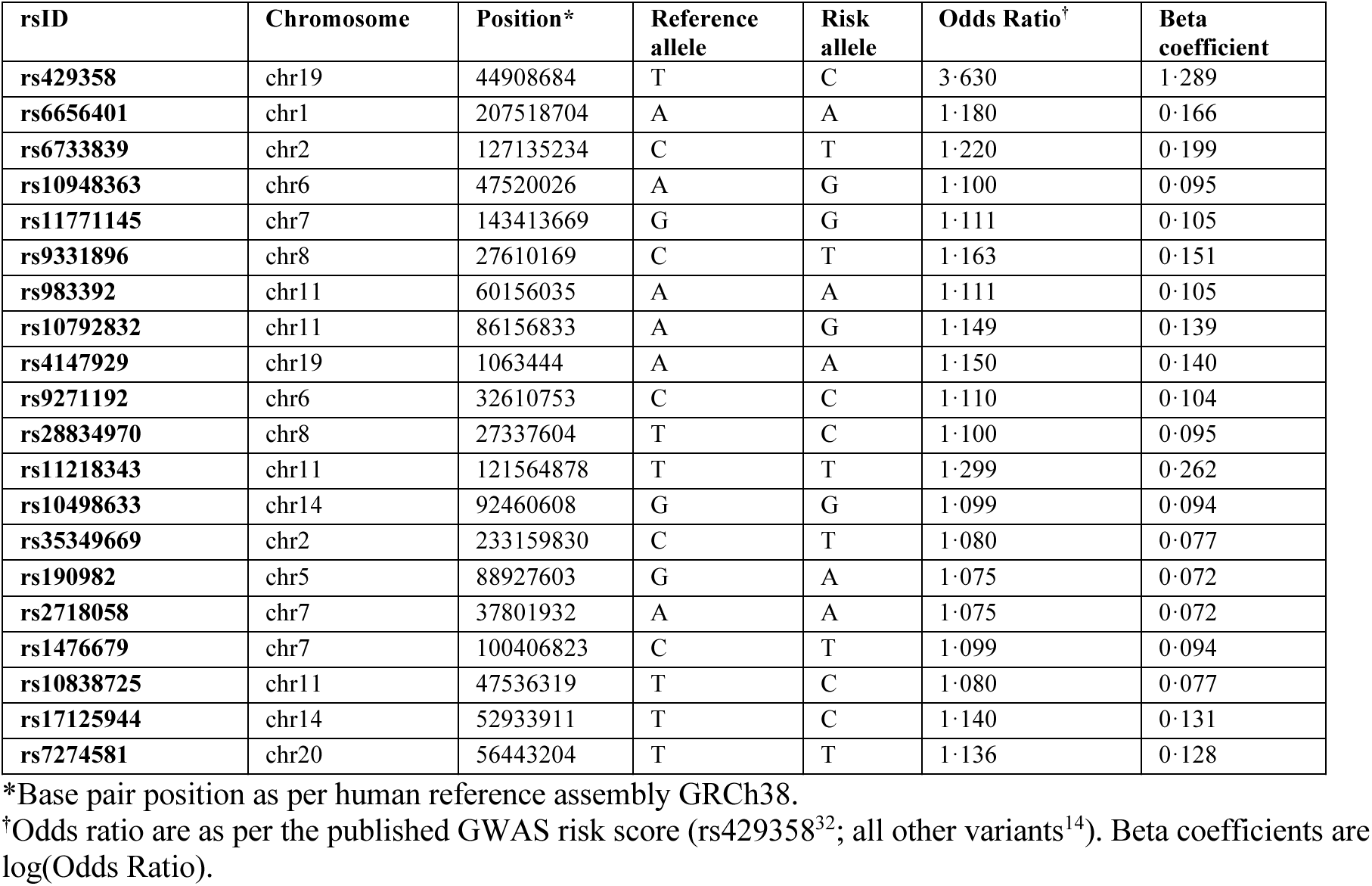
Twenty variants included in polygenic risk model.

**Table S 2.** Comparison of Model Concordance Indices

| Model Description | C-Index | Training Dataset | Data Assessed |
| --- | --- | --- | --- |
| **APOE versus PRS (Instantaneous Age Adjusted Risk)** |  |  |  |
| **APOE Only** | 0.709 | 611 controls/88 cases (ADNI) | 5-fold Cross-validation Same as “Training Data” |
| **PRS Only** | 0.731 | 611 controls/88cases (ADNI) | 5-fold Cross-validation Same as “Training Data” |
| **Training Cohort - MRI only versus MRI/PRS (Age Stratified 1-8 years after baseline Imaging)** |  |  |  |
| **MRI only** | 0.790 | Training Cohort (Described in Table 2) | Same as “Training Data” |
| **MRI/PRS** | 0.820 | Training Cohort (Described in Table 2) | Same as “Training Data” |
| **Validation Cohort - MRI only versus MRI/PRS (Age Stratified 1-8 years after baseline Imaging)** |  |  |  |
| **MRI only** | 0.788 | Training Cohort (Described in Table 2) | Independent Multisite Validation Cohort (Table 2) |
| **MRI/PRS** | 0.807 | Training Cohort (Described in Table 2) | Independent Multisite Validation Cohort (Table 2) |
| **Reverse Training Cohort - MRI only versus MRI/PRS (Age Stratified 1-8 years after baseline Imaging)** |  |  |  |
| **MRI only** | 0.814 | Independent Multisite Validation Cohort (Described in Table 2) | Training Cohort (Described in Table 2) |
| **MRI/PRS** | 0.821 | Independent Multisite Validation Cohort (Described in Table 2) | Training Cohort (Described in Table 2) |
| **Validation Cohort - MRI only versus MRI/PRS (Age Stratified 1-8 years after baseline Imaging)** |  |  |  |
| **MRI only** | 0.779 | Independent Multisite Validation Cohort (Described in Table 2) | Training Cohort (Table 2) |
| **MRI/PRS** | 0.814 | Independent Multisite Validation Cohort (Described in Table 2) | Training Cohort (Table 2) |
| **CSF-derived Amyloid Measures versus MRI only and MRI/PRS (Age Stratified 1-8 years after baseline Imaging)** |  |  |  |
| **CSF** | 0.809 | N = 518 events = 63 Subset of Training Cohort (Described in Table 2) | 5-fold Cross-validation Same as “Training Data” |
| **MRI only** | 0.793 | N = 518 events = 63 Subset of Training Cohort (Described in Table 2) | 5-fold Cross-validation Same as “Training Data” |
| **MRI/PRS** | 0.815 | N = 518 events = 63 Subset of Training Cohort (Described in Table 2) | 5-fold Cross-validation Same as “Training Data” |
| **Study 1: PET-derived Amyloid Measures versus MRI only and MRI/PRS (Age Stratified 1-4 years after baseline Imaging)** |  |  |  |
| **Amyloid-PET (UCB)** | 0.832 | N = 482 events = 39 Subset of Training Cohort (Described in Table 2) | 5-fold Cross-validation Same as “Training Data” |
| **MRI only** | 0.762 | N = 482 events = 39 Subset of Training Cohort (Described in Table 2) | 5-fold Cross-validation Same as “Training Data” |
| **MRI/PRS** | 0.785 | N = 482 events = 39 Subset of Training Cohort (Described in Table 2) | 5-fold Cross-validation Same as “Training Data” |

| Study 2: PET-derived Amyloid Measures versus MRI only and MRI/PRS (Age Stratified 1-3 years after baseline Imaging) |  |  |  |
| --- | --- | --- | --- |
| Amyloid-PET (sPAP) | 0.779 | N = 413 events = 44 Subset of Training Cohort (Described in Table 2) | 5-fold Cross-validation Same as "Training Data" |
| MRI only | 0.792 | N = 413 events = 44 Subset of Training Cohort (Described in Table 2) | 5-fold Cross-validation Same as "Training Data" |
| MRI/PRS | 0.829 | N = 413 events = 44 Subset of Training Cohort (Described in Table 2) | 5-fold Cross-validation Same as "Training Data" |

**Table S 3.** Comparison of Different Genetics Models.

| Dataset | Feature | C-Index | Hazard Ratio | -log10(P) |
| --- | --- | --- | --- | --- |
| NACC | PRS | 0.63 | 1.5 | 5.1 |
| NACC | APOE4 Heterozygous | 0.62 | 2 | 2.6 |
| NACC | APOE4 Homozygous | 0.62 | 5.1 | 4.9 |
| ADNI | PRS | 0.72 | 2.1 | 13.2 |
| ADNI | APOE4 Heterozygous | 0.70 | 4.9 | 9.7 |
| ADNI | APOE4 Homozygous | 0.70 | 8.2 | 8.3 |

**Table S 4.** Survival probability by year, genotype (number of e4 alleles present) and dataset.

| # of e4 | 0 | 0 | 1 | 1 | 2 | 2 |
| --- | --- | --- | --- | --- | --- | --- |
| Dataset | ADNI | NACC | ADNI | NACC | ADNI | NACC |
| Baseline | 1.000 | 1.000 | 1.000 | 1.000 | 1.000 | 1.000 |
| Year 1 | 0.984 | 0.986 | 0.95 | 0.957 | 0.891 | 1.000* |
| Year 2 | 0.966 | 0.961 | 0.829 | 0.896 | 0.766 | 0.804 |
| Year 3 | 0.932 | 0.921 | 0.739 | 0.835 | 0.643 | 0.731 |
| Year 4 | 0.925 | 0.901 | 0.692 | 0.776 | 0.643 | 0.511 |
| Year 5 | 0.91 | 0.871 | 0.63 | 0.75 | 0.536 | 0.365 |
| Year 6 | 0.91 | 0.855 | 0.581 | 0.716 | 0.268 | 0.365 |
| Year 7 | 0.874 | 0.834 | 0.528 | 0.697 | 0.268 | 0.365 |
| Year 8 | 0.825 | 0.78 | 0.403 | 0.636 | 0.268 | 0.365 |

\*Due to the size of the dataset, there were no individuals homozygous for APOE e4 who progressed to dementia in the first year in the validation cohort.

**Figure S 1.**
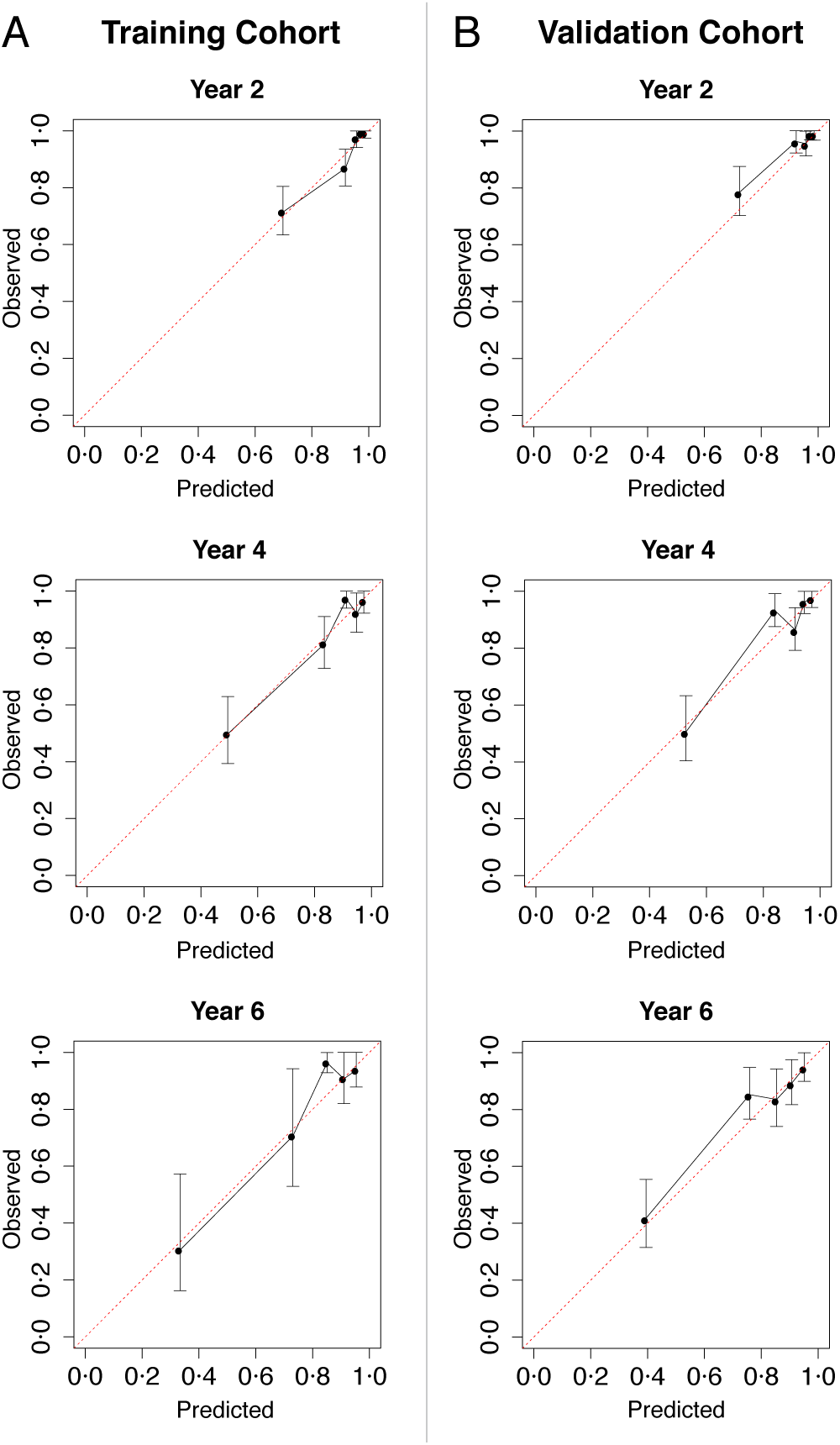
Calibration plots in the training and validation cohorts for prediction 2, 4, & 6 years prior to a progression event.

**Figure S 2.**
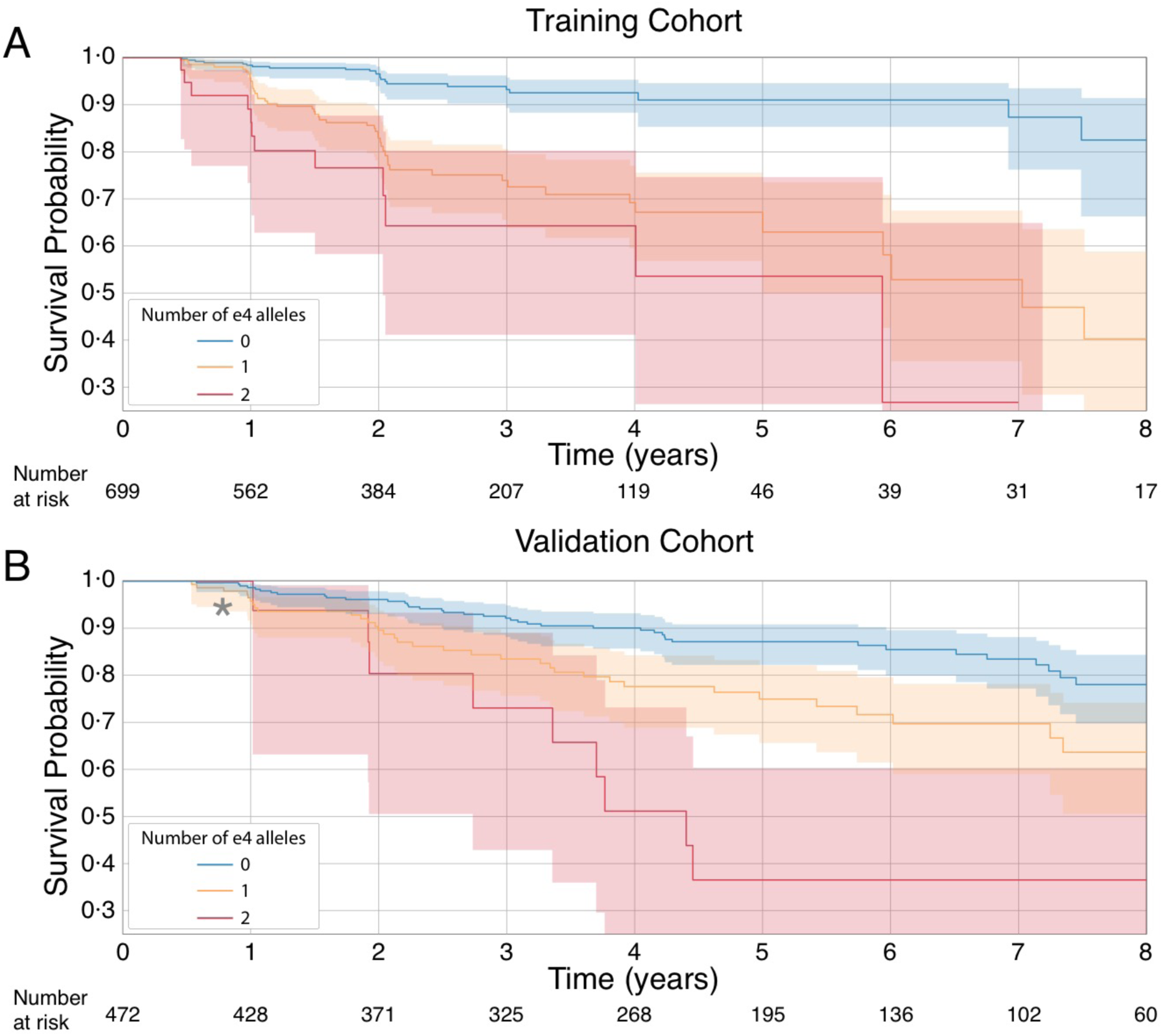
Plots illustrating the survival probability of individuals categorized by APOE genotype. Due to small sample size in the validation cohort at year 1. **\***There were no individuals homozygous for APOE e4 who progressed to dementia in the first year in the validation cohort.

